# The composition of the microbiota in the full-term fetal gut and amniotic fluid: a bovine caesarean section study

**DOI:** 10.1101/2020.09.28.309476

**Authors:** Aleksi Husso, Leen Lietaer, Tiina Pessa-Morikawa, Thomas Grönthal, Jan Govaere, Ann Van Soom, Antti Iivanainen, Geert Opsomer, Mikael Niku

## Abstract

The fetal development of the intestinal immune system is stimulated by the maternal microbiota, but it is still unclear whether viable bacteria exist in the healthy fetus. Analysis of such low microbial biomass environments are challenging due to contamination issues. The aims of the current study were to assess the bacterial load and characterize the bacterial composition of the amniotic fluid and meconium of full-term calves, leading to a better knowledge of prenatal bacterial seeding of the fetal intestine. Amniotic fluid and rectal meconium samples were collected during and immediately after elective caesarean section, performed in 25 Belgian Blue cow-calf couples. The samples were analyzed by qPCR, bacterial culture using GAM agar and 16S rRNA gene amplicon sequencing. To minimize the effects of contaminants, we included multiple technical controls and stringently filtered the 16S rRNA gene sequencing data to exclude putative contaminant sequences. The meconium samples contained a significantly higher amount of bacterial DNA than the negative controls and 5 of 24 samples contained culturable bacteria. In the amniotic fluid, the amount of bacterial DNA was not significantly different from the negative controls and all samples were culture negative. Bacterial sequences were identified in both sample types and were primarily of phyla Proteobacteria, Firmicutes, Bacteroidetes and Actinobacteria, with some individual variation. We conclude that most calves encounter *in utero* maternal-fetal transmission of bacterial DNA, but the amount of bacterial DNA is low and viable bacteria are rare.

## Introduction

Characterizing the very first intestinal bacteria is essential for a better understanding of the co-development of the newborn and its intestinal microbiome. Host-microbiome interactions enable early-life education and maturation of the immune system during a *window of opportunity*. The use of prenatal and intrapartum antibiotics may disturb this process (Dierikx et al., 2020). However, the timing of the microbial colonization of the mammalian gut is still unclear (Perez-Munoz et al., 2017; Korpela and de Vos, 2018; Liu et al., 2019; Guzman et al., 2020). Vertical as well as environmental transmission of bacteria occur during and after birth and seed the neonatal gastrointestinal (GI) tract (Funkhouser and Bordenstein, 2013; Korpela and de Vos, 2018). Transmission of an orally inoculated *Enterococcus faecium* strain in pregnant mice to the meconium of their fetuses has been described (Jimenez et al., 2008).

Recent studies in humans, mice and cattle have reported the identification of microbial DNA or culturable bacteria in the mammalian fetal environment (Aagaard et al., 2014; Severovic et al., 2019; Al Alam et al., 2020; Rackaityte et al., 2020; Guzman et al., 2020). Nevertheless, others interpret such observations as intrauterine infections or contamination (Lauder et al., 2016; Eisenhofer et al., 2018; de Goffau et al., 2019; Hockney et al., 2020; Olomu et al., 2020; Theis et al., 2020a, 2020b). The newborn’s GI tract is a low microbial biomass environment, implying multiple challenges for performing reliable microbiome analyses (Glassing et al., 2016; Eisenhofer et al., 2018; Stinson et al., 2018). Besides the technical limitations inherent to DNA sequencing, there are practical and ethical concerns while collecting samples in humans. First-pass meconium has been used as a proxy to assess the fetal intestinal microbiome, although samples are often collected hours after birth and even breastfeeding, suggesting potential postnatal effects on the microbial composition of the meconium (Collado et al., 2016; Liu et al., 2019; Stinson et al., 2019). Mammalian animal models can overcome several of these ethical and practical difficulties, providing insights in the fetal gastrointestinal microbiome.

In the current study, the microbial composition of the full-term fetal gut and the corresponding amniotic fluid was assessed in Belgian Blue cow-calf couples. In this double-muscled beef breed, caesarean sections (C-sections) are performed on a routine basis during the very early stages of parturition, while fetal membranes are still intact, rendering this breed a highly suitable model for full-term gestation microbiome studies.

Besides the potential role of the cow model for research, there is also a significant interest in the composition and the development of the calf’s intestinal microbiome. Gut health and growth performance during the first weeks of life are main drivers for cost effective livestock rearing (Thornton Philip, 2010; Lorenz et al., 2011). Few studies have been conducted on the intestinal microbiome in vaginally born neonatal calves. These have reported low numbers of bacteria, mostly belonging to the phyla Proteobacteria, Firmicutes, Actinobacteria and Bacteroidetes (Alipour et al., 2018; Klein-Jöbstl et al., 2019; Guzman et al., 2020). Shared microbiota were found between calf meconium and the maternal vaginal vestibulum (Alipour et al., 2018; Yeoman et al., 2018; Klein-Jöbstl et al., 2019). To the best of our knowledge, no microbiome studies have yet been performed on calves born by C-section, and no studies are available to assess the association between the microbiome of the calf’s meconium and the microbiome of the corresponding amniotic fluid.

Our aims were to assess the bacterial load and characterize the bacterial composition of the amniotic fluid and meconium of full-term neonatal calves. To this end, samples were collected during elective C-section, and analyzed by qPCR, bacterial culture under carefully controlled and validated conditions, and 16S rRNA gene amplicon sequencing. To minimize reagent and environmental bacterial DNA contaminants, we included multiple technical controls and stringently filtered the 16S rRNA gene sequencing data to exclude putative contaminant sequences.

## Materials and methods

All experimental procedures were approved by the institutional ethics and animal welfare committee of the Faculty of Veterinary Medicine (EC2018/002 - Ghent University, Belgium). The cows’ owners were informed about the study and gave their written consent.

### Study design

The sampling of this study was performed at the teaching hospital of the Department of Reproduction, Obstetrics, and Herd Health of the Faculty of Veterinary Medicine in Ghent (Belgium), where pregnant Belgian Blue beef cows of different herds (parity between 1 and 5) were housed for an elective C-section.

In total 25 Belgian Blue cows and their calves (7 males and 18 females) were sampled from November 2017 until March 2019. The cows were housed in tie-stalls at the facility for 9.5 d ± 5.8 (mean ± standard deviation) prior to C-section and had ad libitum access to hay and water. During this period, rectal temperature was measured twice daily, and calving indicators such as udder distension, teat filling, pelvic ligament relaxation, vaginal discharge, vulvar edema, and behavioral changes were monitored every two hours by graduate veterinary students.

Prior to elective C-section, in cows that exhibited a drop in temperature, cervical dilation was assessed by manual palpation. The vulvar region was cleaned with iodine soap and water. A gloved hand was inserted vaginally and the opening of the portio vaginalis cervicis was estimated. For the present study, elective C-section was performed when the cow had a minimal cervical dilation of eight centimeters, with no rupture of the fetal membranes prior to surgery. All cows were healthy according to their vital parameters (heart rate, temperature, respiratory rate) and there was no clinical evidence of intrauterine infection or contamination.

### Sampling

Prior to surgery, the cows were restrained in a standing position in a surgery crush specifically designed for cattle. C-section procedure was done as described by Kolkman et al. (2007). Briefly, the surgical area (left flank) was washed and disinfected, an abdominal incision was made, and part of the uterus was exteriorized for uterotomy. The allantoic sac was opened up to expose the intact amniotic sac. Amniotic fluid was aspirated through the amniotic membrane, using a sterile 16 G needle (Agani, Terumo Europe, Hamburg, Germany) and sterile 20 ml syringe (B. Braun, Melsungen, Germany). Within one hour after sampling, the retrieved volume was aliquoted, under a laminar-flow hood, into 2 sterile 15 ml tubes (188271, Cellstar, Greiner bio-one, Frickenhausen, Germany). In the first tube, 6 ml of amniotic fluid was dissolved in 3 ml of glycerol (≥99 %, G2025, Sigma-Aldrich (Merck), Overijse, Belgium) and subsequently stored at −80 °C to be used in culture experiments. In the second tube, 12 ml of amniotic fluid was stored at −80 °C to be used for bacterial DNA extraction.

Meconium samples were acquired directly from the calves’ rectum, immediately after birth (no more than 30 minutes). Until the moment of sampling, calves laid on a clean concrete floor, with access to neither the dam, nor colostrum. The perineum of the calf was dried with a clean paper towel and disinfected with 70 % ethanol. A sterile double-guarded equine uterine culture swab (Har-vet, 17705, Spring Valley, USA; or Minitube, 17214/2950, Tiefenbach, Germany) was gently introduced in the rectum and the swab was exposed. Samples were taken in duplicate, one stored immediately at −80 °C with no additives, and the other stored at −80 °C in a sterile 2 ml cryovial containing 1 ml of a 30 % glycerol solution, prepared by diluting glycerol (≥99 %, G2025, Sigma-Aldrich (Merck), Overijse, Belgium), in ultra-pure, nuclease-free water (W4502, Sigma-Aldrich (Merck), Overijse, Belgium) to a final 30 % concentration.

Negative field controls were processed in the surgery room, using the same sampling procedures and disposables. In total, 16 empty, sterile double-guarded equine uterine culture swabs (10 Har-vet swabs and 6 Minitube swabs) were included for the meconium sampling, 5 of the Har-vet swabs stored in 1 ml of a 30 % glycerol stock solution, the others with no additives. Additionally, 12 negative controls were included for the amniotic fluid sampling, aspirating ultra-pure, nuclease-free water (W4502, Sigma-Aldrich (Merck), Overijse, Belgium) instead of amniotic fluid. For 8 of them, the ultra-pure, nuclease-free water was stored with no additives, and for 4 of them, 6 ml ultra-pure, nuclease-free water was dissolved in 3 ml of glycerol.

All samples were shipped on dry ice to the laboratory of the Department of Veterinary Biosciences of the Faculty of Veterinary Medicine in Helsinki (Finland) for further processing.

### Culture

#### Validation of culture media

One of our aims was to define a single bacterial culture medium, capable of sustaining a majority of the calf intestinal core microbiota. Consequently, this single medium could be used for culturing the meconium and amniotic fluid samples, avoiding further splitting or dilution of the low biomass samples. We first tested GAM “Nissui” (Gifu Anaerobic Medium Agar, Code 05420, HyServe, Germany), YCFA (Yeast extract, Casitone and Fatty Acid), LB (Lysogeny Broth, BD) and BB (Trypticase Soy Agar supplemented with Bovine Blood 211043, Tammer BioLab Oy, Tampere, Finland) media for the ability to sustain calf intestinal core microbiota, by plating each type of medium anaerobically with feces of a healthy 7 day old calf (Lopez-Siles et al., 2012; Alipour et al., 2018). After 7 days of growth, mixed cultures were extracted from the plates, 16S rRNA gene amplicon sequenced, and compared to the core microbiota previously observed in fecal samples of young calves (Alipour et al., 2018).

#### Sample culturing

Bacterial culture on GAM-agar plates was performed for 24 meconium samples (5 corresponding negative controls) and 24 amnion samples (4 negative controls). The frozen samples were first transferred to an anaerobic workstation (Ruskinn Concept Plus), mixed thoroughly and plated, using an aseptic technique, at +37 °C. After this, the samples were transferred to a laminar flow cabinet and plated in aerobic conditions at +37 °C. The culture plates were checked for growth daily for 14 days. All visible bacterial colonies were subcultured until pure isolates were obtained. Fresh cultures were then identified using MALDI-TOF (Bruker Microflex LT) at the Central laboratory of the Faculty of Veterinary Medicine (Helsinki, Finland).

Samples were prepared using MALDI Biotyper MSP Identification Standard Method v 1.1. Mass spectra were analyzed in a mass/charge range from 2000 to 20 000 Da with MBT Compass v4.1 on flexControl v3.4 (Bruker Daltonik GmbH) using BDAL-7311 as the reference library. The Bruker Bacterial Test Standard (RUO) (Bruker Daltonik GmbH) was used for instrument calibration. If the identification confidence score was <2.00, further identification was done with 16S rRNA gene amplicon Sanger sequencing at the Institute of Biotechnology (University of Helsinki, Finland).

### DNA extraction

DNA from the meconium samples (N = 25) and corresponding negative controls (N = 11) were extracted using ZymoBIOMICS DNA Miniprep Kit (Zymo Research, Irvine, CA, USA) according to the manufacturer’s instructions, with minor modifications to the protocol as described previously (Alipour et al., 2018; Husso et al., 2020). ZymoBIOMICS™ Microbial Community Standard and an in-house fecal standard were processed with the meconium samples and no-template-controls were included in every batch.

For the amniotic fluid samples (N = 23) and the corresponding negative controls (N = 8), 2 ml of each sample was first centrifuged in a microcentrifuge at 16100 × g 10 min in +4 °C. Most of the supernatant was removed and 750 μl of ZymoBIOMICS Lysis Solution and 19 μl of proteinase K (D3001-2-20/D3001-2-5, Zymo Research) were added to the remaining 200 μl of the samples and incubated for 30 min at +55 °C. After these steps, the same protocol as applied for the meconium samples was followed.

All manipulations of the tubes during the process were performed in a laminar flow cabinet, and the workplace, instruments and pipettes were cleaned routinely with 10 % bleach. Certified DNA, RNase, DNase and PCR inhibitor free tubes (STARLAB International, Germany) and Nuclease-free Water (Ambion, Thermo Fisher Scientific, USA) were used for DNA extraction and downstream analyses. All extracted DNA was stored at −80 °C.

### Quantitative PCR

The bacterial 16S rRNA gene copy numbers in the meconium, amnion and negative control samples were determined using quantitative PCR. The analyses were performed as described previously (Alipour et al., 2018; Husso et al., 2020), with the exception that the PCR master mix was first treated using a dsDNAse based decontamination kit (Enzo Life Sciences, Farmingdale, New York). The amplification was performed using the Bio-Rad CFX96 instrument (Bio-Rad, Hercules, California) and universal eubacteria probe and primers (Nadkarni et al., 2002). A standard series and negative controls were included in every run. The data were analyzed using the Bio-Rad CFX Maestro software. The results were normalized to the averages of absolute 16S rRNA gene copy numbers in the negative controls included in each run.

### Library preparation and 16S rRNA gene amplicon sequencing

The V3-V4 region of the 16S rRNA gene amplicons was sequenced using the Illumina MiSeq platform in the DNA core facility of the University of Helsinki, as described previously (Alipour et al., 2018; Husso et al., 2020). In total, 23 amniotic fluid samples and 23 meconium samples of the same cow-calf couple were sequenced.

The ZymoBIOMICS Microbial Community Standard (Zymo Research, USA) and the in-house adult cow fecal standard were pre-amplified with 12 cycles and all other sample types, including negative controls, with 21 cycles.

The observed composition and abundances for the commercial standard matched the expected composition provided by the manufacturer (data not shown).

### Bioinformatics

The detailed bioinformatics pipeline is described in the Supplementary materials. Briefly, the read quality was first inspected with FastQC and MultiQC (Andrews, 2011; Ewels et al., 2016). Leftover primes and spacers were then trimmed with Cutadapt v1.10 (Martin, 2011). A mapping file was created for QIIME2 and validated with Keemei (Rideout et al., 2016). The FASTQ-files were imported to QIIME2 v2019.4, where the DADA2 plugin was used to denoise and quality filter the reads, call ASVs and generate a feature table (Callahan et al., 2016; Bolyen et al., 2019). A naïve Bayes classifier was trained in QIIME2 against SILVA v132 99 % database, extracted to only include the V3-V4 region and used to assign taxonomy to ASVs (Quast et al., 2013; Bokulich et al., 2018). Sequences derived from chloroplasts or mitochondria were removed and singletons were filtered out, leaving only bacteria with at least phylum-level identification.

The processed data was *in silico* filtered to remove ASVs which represented probable contaminants, as described previously (Husso et al., 2020). Briefly, an ASV was removed if its prevalence in actual samples was ≤2× its prevalence in instrument controls (DNA extracted from empty sampling instruments only handled aseptically in a laminar flood cabinet) or its prevalence in field controls (empty sampling instruments exposed to the surgery room environment), and if its mean relative abundance in actual samples was ≤10× its mean abundance in instrument controls or its mean abundance in field controls. The filtering was performed separately for meconium and amnion samples. If less than 500 reads remained after the decontamination, as was the case for six meconium samples, the sample was removed from further analyses.

### Statistics

The 16S rRNA gene qPCR results from samples and negative controls were compared using two-tailed Mann-Whitney U test in IBM SPSS Statistics 25.

Shannon diversity indices were calculated for genus-level data with the R package phyloseq (McMurdie and Holmes, 2013). The Kruskal-Wallis rank sum test was used to compare the bacterial community structure of both sample types in RStudio (R Studio Team, 2020).

The PCoA figures were plotted using ASV and genus-level data and Bray-Curtis distances with the R package phyloseq (McMurdie and Holmes, 2013). Permutational analysis of variance (PERMANOVA) and multivariate homogeneity of group dispersions (betadisper) were calculated using the R package vegan, using 9999 permutations (Oksanen et al., 2019).

DESeq2 was used to explore differentially abundant ASVS by calculating differential expression between sample groups (Love et al., 2014).

The LEfSe (Linear discriminant analysis Effect Size) web application was used to identify the taxons most likely explaining the differences between the sample types (Segata et al., 2011).

An ecologically organized heatmap of the top 40 most abundant ASVs was created with the R package phyloseq (Rajaram and Oono, 2010; McMurdie and Holmes, 2013).

Spearman correlations and their Bonferroni corrected p-values were calculated using R-package Hmisc 4.3.0 (Harrel, 2019).

Differences were considered significant at P < 0.05.

## Results

### Quantification of bacterial 16S rRNA in amniotic fluid and meconium samples by qPCR

The 16S rRNA gene copy numbers in amniotic fluid and meconium samples were assessed by qPCR (Fig. 1). To minimize variation between qPCR runs, the copy numbers were normalized to the averages of the negative controls in the same run as explained in Materials and Methods. In the amniotic fluid samples, there was no significant difference (P = 0.104) between the normalized 16S rRNA gene copy numbers of the samples (N = 23; mean = 2.4; SD = 2.1) and the negative controls (N = 8; mean = 1; SD = 0.53).

**Figure 1.**
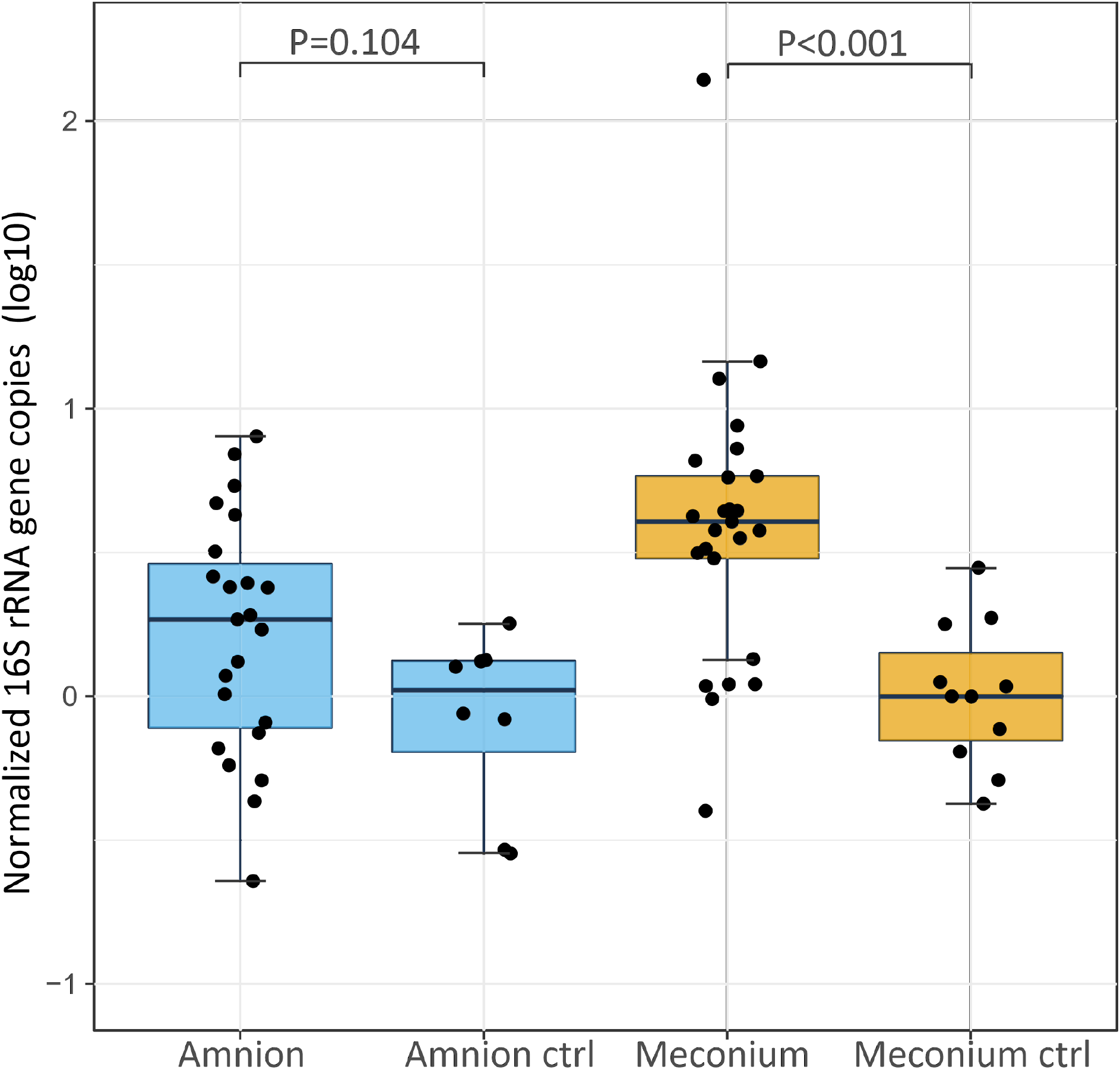
16S rRNA gene copy numbers (fold change vs. negative controls) for amniotic fluid (n = 23, negative controls n = 8) and meconium samples (n = 25, negative controls n = 11). Samples were collected during elective C-section in Belgian Blue cow-calf pairs. The copy number of each sample was normalized to the average copy number of the corresponding negative controls and log10 transformed. The midline of the box is the median, with the upper and lower limits of the box being the third and first quartile. The whiskers extend up to 1.5 times the interquartile range from the box to the furthest data point within that distance.

In the meconium samples, the 16S rRNA gene copy number (N = 25; mean = 9.9; SD = 27) was significantly higher (P < 0.001) than in the negative controls (N = 11; mean = 1; SD = 0.86).

### Decontamination of the 16S rRNA gene amplicon sequencing data

We performed a stringent filtering of the 16S rRNA gene sequencing data to remove the amplicon sequence variants (ASVs) potentially stemming from reagent contaminants and environmental bacterial DNA (Table 1).

**Table 1.** Mean read counts (standard deviation) detected in meconium and amniotic fluid samples and their negative controls. Only sequences with at least phylum-level identification were retained for further analyses. If less than 500 reads remained after in silico decontamination, as in six meconium samples, the sample was removed from further analyses.

| Sample type | Raw |  | Processed |  | Decontaminated |  |
| --- | --- | --- | --- | --- | --- | --- |
|  | N | Reads | N | Reads | N | Reads |
| Meconium | 23 | 92266<br>(36235) | 23 | 57082<br>(25543) | 17 | 4189<br>(8931) |
| Meconium control | 11 | 126800<br>(21164) | 11 | 84023<br>(14585) | - | - |
| Amnion | 23 | 127797<br>(22376) | 23 | 83279<br>(15120) | 23 | 3338<br>(2922) |
| Amnion control | 6 | 136488<br>(12663) | 6 | 90802<br>(8998) | - | - |

The decontamination was based on the prevalence and relative abundance of each ASV in the samples and negative controls (see Materials and Methods).

### Microbial composition of amniotic fluid and meconium samples

The microbiota of both sample types consisted primarily of Proteobacteria, Firmicutes, Bacteroidetes and Actinobacteria phyla, with individual variation (Fig. 2). In both sample types, the inter-individual variation increased at lower taxonomic levels (Supplementary Table 1 and Fig. 3). The most abundant bacterial genera in meconium were *Delftia*, *Staphylococcus* and *Clostridium sensu stricto 1*, while the most prevalent genera were *Delftia*, *Acinetobacter*, unclassified *Burkholderiaceae*, *Staphylococcus* and *Corynebacterium 1* (Supplementary Table 1). In the amniotic fluid samples, *Staphylococcus*, *Streptococcus, Delftia, Sphingomonas* and *Enterococcus* were the most abundant genera, and *Delftia*, *Streptococcus*, *Staphylococcus, Sphingomonas* and *Acinetobacter* were the most prevalent genera (Supplementary Table 1). The observed composition and abundances for the commercial standard matched the expected composition provided by the manufacturer.

**Figure 2.**
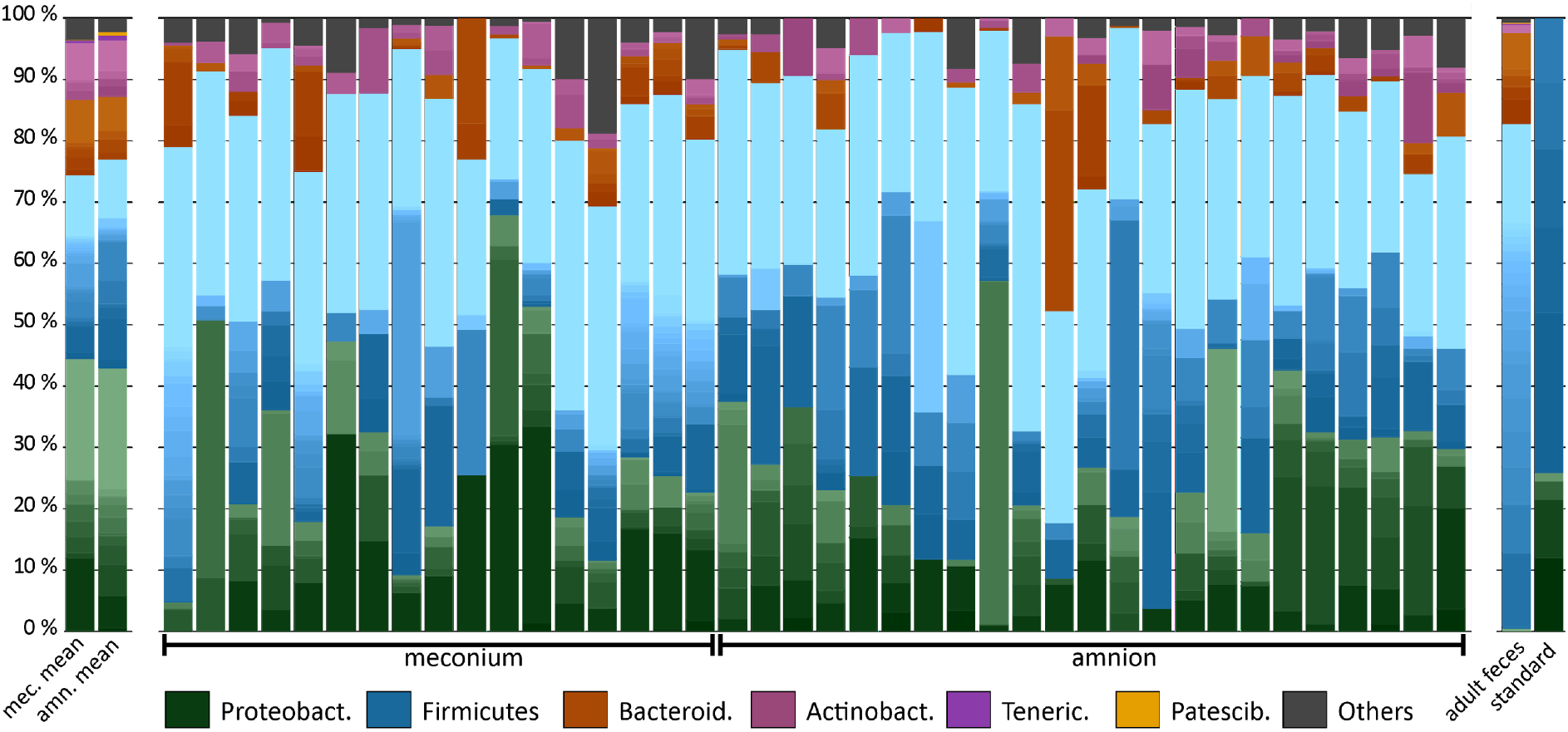
Microbiota composition in bovine meconium and amniotic fluid samples collected during elective C-section, adult feces, and a commercial community composition standard. The main colors indicate the bacterial phyla. Within phyla, the shades indicate bacterial genera. The lightest shade of each phylum shows the combined abundance of the least abundant genera (with a maximum of <0.5% of total).

A heatmap analysis of the 40 most abundant ASVs shows the difference between the sample types in more detail (Fig. 3).

**Figure 3.**
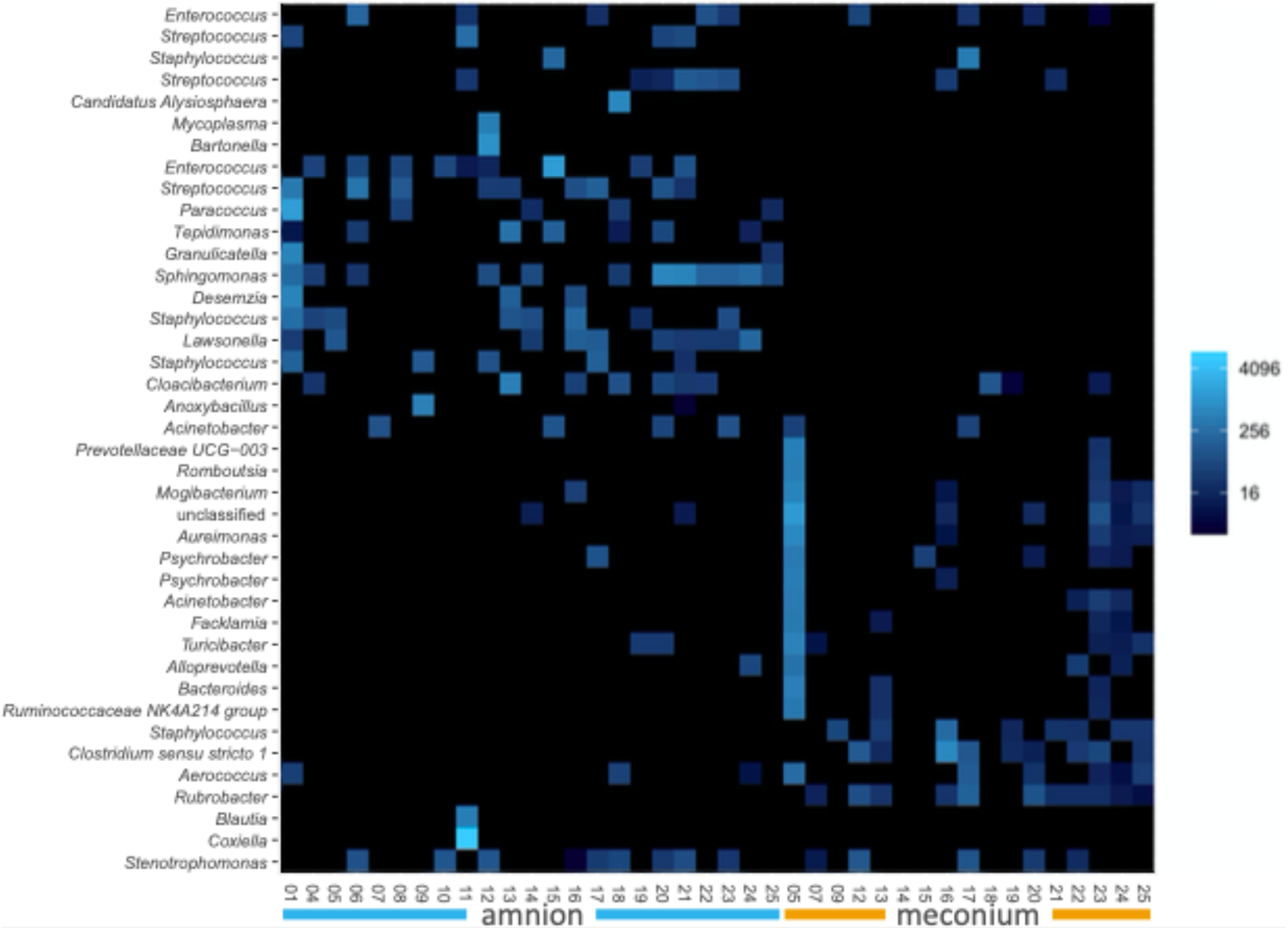
Ecologically organized heatmap (NMDS, Bray) of the 40 most abundant ASVs in bovine meconium and amniotic fluid samples collected during elective C-section, sorted by sample type. The taxon names are presented as genus level identifications. Color scale indicates relative abundance and is a log transformation with base 4.

The alpha diversity (Shannon index) at the genus level was not significantly different between the meconium and amniotic fluid samples (Fig. 4, P = 0.889).

**Figure 4.**
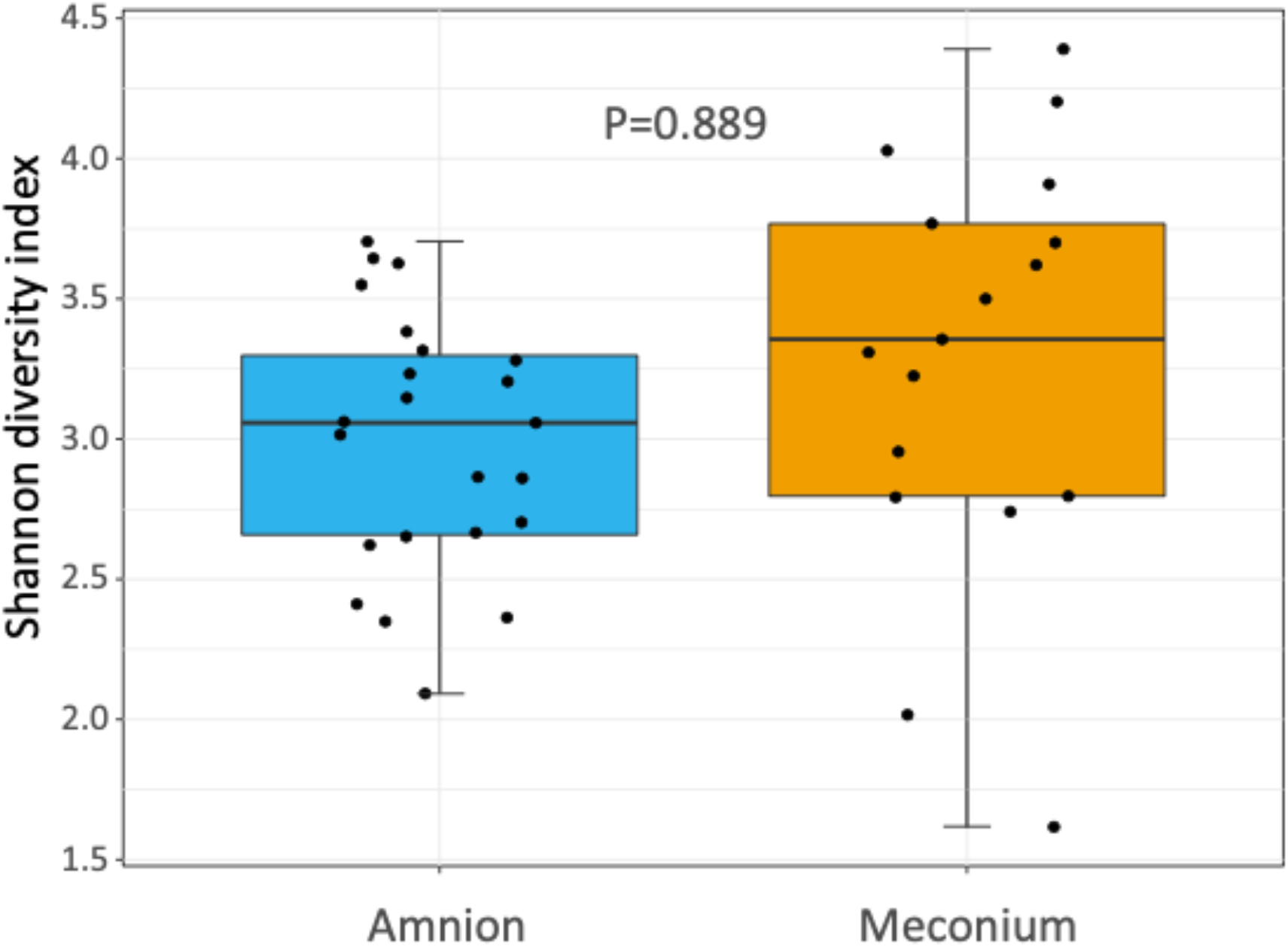
Shannon diversity index showing the bacterial community structure (at genus level) of amniotic fluid and meconium samples, collected during elective C-section in Belgian Blue cow-calf pairs. Boxplot as in Fig 1.

### Comparison of amniotic fluid and meconium microbiota composition

The microbiota composition was significantly different for amniotic fluid and meconium samples at the ASV (PERMANOVA, P < 0.001, R2 = 0.049, Fig. 5 A), and genus level (PERMANOVA, P < 0.001, R2 = 0.062, Fig. 5 B).

**Figure 5.**
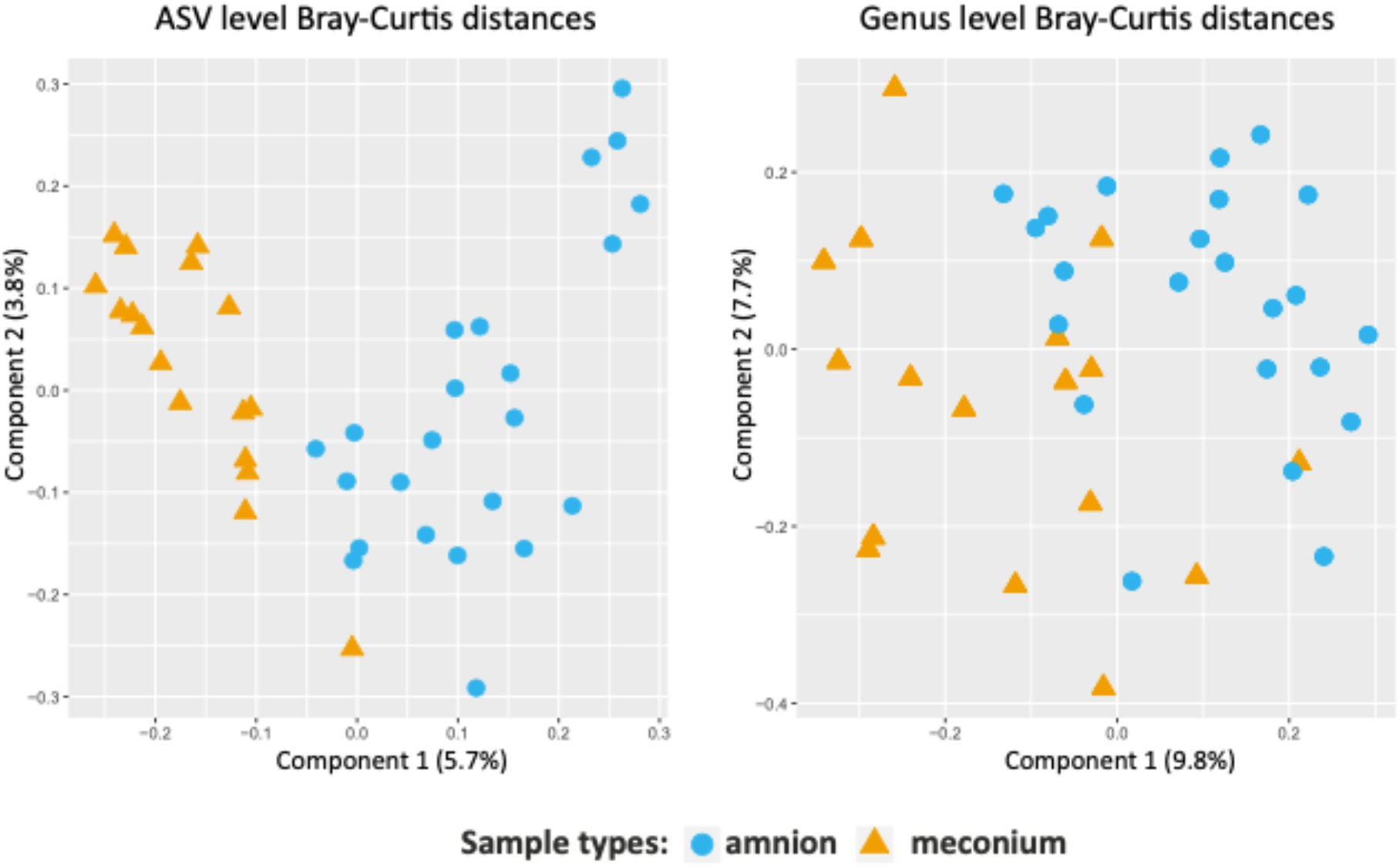
Comparison of the bacterial community structure of amniotic fluid and meconium. 5 A: PCoA on Bray-Curtis distances based on ASV level data. Figure 5 B: PCoA on Bray-Curtis distances based on genus level data. Colors and shapes indicate the sample types.

According to the DESeq2 analysis, three ASVs were more abundant in meconium than in amniotic fluid: one *Staphylococcus* ASV (log2 Fold Change = 25.084, P_adj_ < 0.001), one *Rubrobacter* ASV (log2 Fold Change = 8.337, P_adj_ < 0.001), and one *Clostridium* ASV (log2 Fold Change = 8.055, P_adj_ = 0.004).

In amniotic fluid, one *Sphingomonas* ASV (log2 Fold Change = −8.420, P_adj_ < 0.001) and one *Staphylococcus* ASV (log2 Fold Change = −23.438, P_adj_ < 0.001) were more abundant than in meconium.

By LefSe (Linear discriminant analysis effect size) analysis including all taxonomic levels, meconium samples had a greater relative abundance of *Clostridiales* and *Rubrobacter* (LDA score (log10) > 4.0) than the amniotic fluid, but a lower relative abundance of Bacillales and *Corynebacteriales* (LDA score (log10) < −4.0).

### Correlations between meconium and amniotic fluid from the same animal

We calculated Spearman rank correlations for the microbiota between meconium and amniotic fluid samples, taking into account ASVs that were present >100 times in all of the samples. Average correlations (ρ) at ASV (N = 17, ρ_avg_ = −0.0201, P_avg_ < 0.05) and genus level (N = 17, ρ_avg_ = 0.1685, P_avg_ < 0.05) were very weak and biologically not significant. All p-values were Bonferroni corrected for multiple testing.

### Bacterial cultures

Based on our testing of multiple media, the GAM (Gifu Anaerobic Medium) agar was selected to assess if there were live bacteria in the samples stored in glycerol (see Materials and Methods), since it was able to sustain the largest number of bacterial core genera found in calf feces, including some additional unique genera not found on any other plate type (Alipour et al., 2018).

All amniotic samples and their negative controls were culture negative, both in anaerobic and aerobic conditions. Five out of 24 meconium samples and 1 out of 5 negative controls were culture positive on GAM agar. The results for each sample are presented in detail in Table 2. Both gram negative and gram-positive strains were isolated from the samples.

**Table 2.**
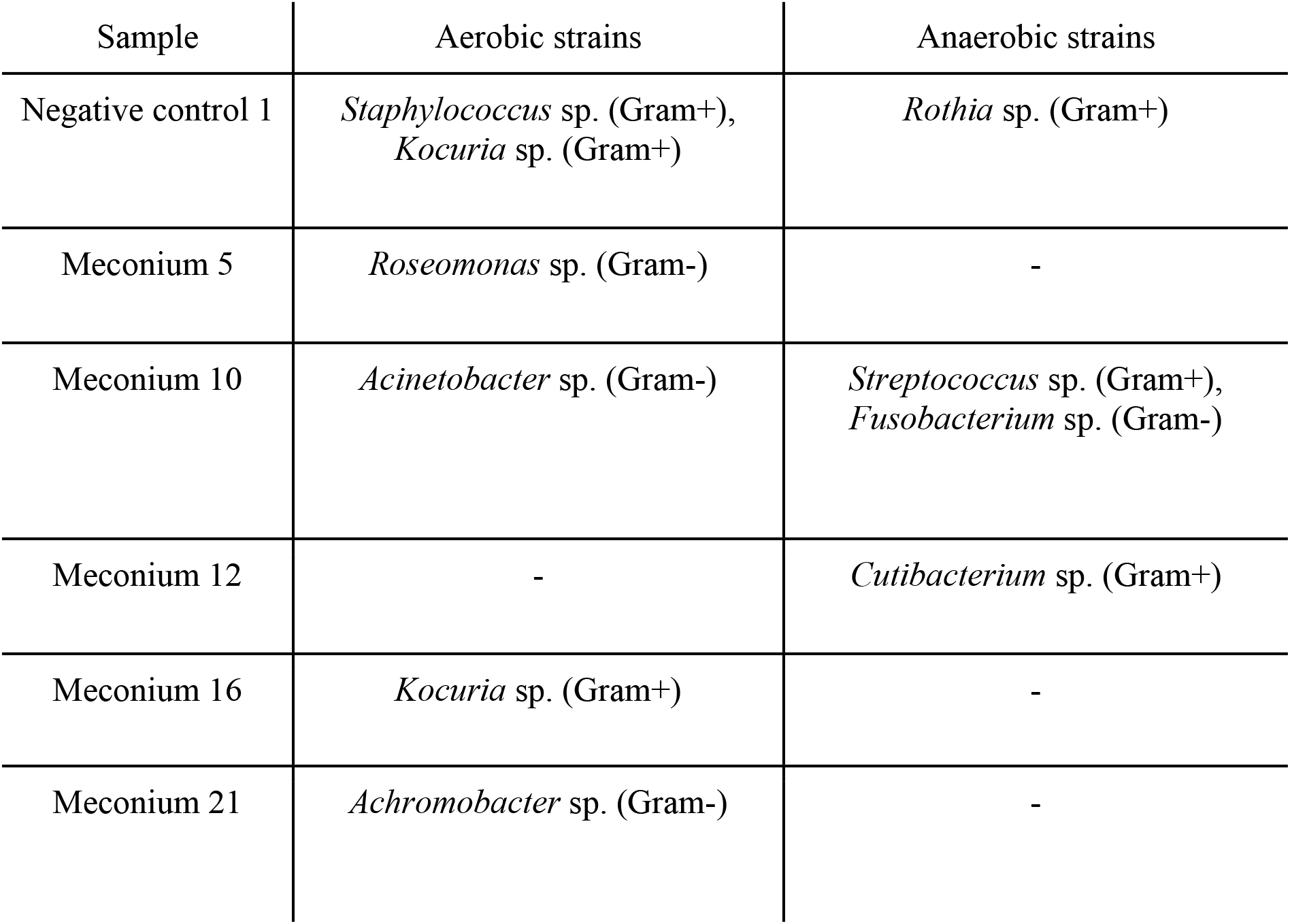
Culture positive meconium samples and identified bacteria. If the identification confidence score was <2.00 for MALDI-TOF, further identification was done with 16S rRNA gene amplicon Sanger sequencing to achieve genus level identification for all isolates.

The Sanger sequence from the *Roseomonas* sp. isolate acquired from a meconium sample was a 100 % identical match to the 16S rRNA gene amplicon sequencing data from the same sample. In addition, the Sanger sequences of *Staphylococcus* sp., *Kocuria* sp., *Roseomonas* sp., and *Acinetobacter* sp. isolates matched the 16S rRNA gene amplicon sequencing data when compared to all meconium samples.

Three bacterial genera could be identified in the negative controls: *Rothia* sp., *Staphylococcus* sp. and *Kocuria* sp.. Of these, only *Kocuria* sp. was also identified from the actual meconium samples.

## Discussion

Vertical transmission of microbes during, and perhaps already before birth is a universal phenomenon across the whole animal kingdom and plays an important role in shaping host immunity and fitness (Funkhouser and Bordenstein, 2013; Vandenplas et al., 2020). Despite the advances in our understanding of the assembly and lifelong effects of the perinatal microbiota, the *in utero* exposure of the fetus to maternal bacteria and bacterial components remains poorly understood (Perez-Munoz et al., 2017; Korpela and de Vos, 2018; Liu et al., 2019).

In the present study, we sampled amniotic fluid during, and the corresponding calves immediately after they were born by elective C-section. The elective C-section is a routine procedure in Belgian Blue beef cattle, since the pelvis of the dam does not allow vaginal birth of the relatively large double-muscled calf (Kolkman et al., 2007). We performed the surgery during the very early stages of parturition, when both fetal membranes (amniotic and allantochorionic membrane) were still intact, allowing us to sample the full-term fetus and the intrauterine environment without exposure to the vaginal microbes and minimal exposure to the environmental microbiota. Meconium samples were collected rectally. The relatively large size of the calves permits the insertion of a double-guarded sampling device into the rectum, minimizing the risk of contamination.

In contrast, in most human studies first-pass meconium was collected from diapers (Hansen et al., 2015; Collado et al., 2016; Liu et al., 2019; Stinson et al., 2019). Because the first-pass meconium is often collected several hours after birth, microbes from the environment and the mother (especially during breastfeeding) have time to colonize the intestine. Hence, this sampling procedure should be seriously questioned for researching the eventual intrauterine transmission of bacteria to the neonatal gastrointestinal tract (Dornelles et al., 2020). In the current study, amniotic fluid samples were collected *in vivo*, while the amniotic membrane was still intact, as has been done in humans during pre-labor C-section (Collado et al., 2016).

Bacterial culture is traditionally used to determine the viable population in a bacterial community. However, standard culture-dependent techniques are known to be inadequate to identify all bacteria. Capturing a broad bacterial diversity would require the use of many different selective media and multiple specific growing conditions (Lau et al., 2016; Lagier et al., 2018). In the current study, one of our aims was to define a single bacterial culture medium, capable of sustaining the majority of the calf intestinal core microbiota. We first tested four different culture media: GAM “Nissui” (Gifu Anaerobic Medium Agar), mostly used for anaerobic bacteria, YCFA (Yeast extract, Casitone and Fatty Acid), containing volatile fatty acids typically supporting the growth of several gut bacteria, LB (Lysogeny Broth) as a general media, and BB (Trypticase Soy Agar supplemented with Bovine Blood) supporting fastidious bacteria by blood enrichment. The GAM medium was found optimal for both aerobic and anaerobic culturing.

Bacteria were successfully cultured from 5 out of 24 meconium samples, representing both gram-negative and gram-positive bacterial genera, known to live in mammalian hosts. *Roseomonas* and *Achromobacter* species are known to act as opportunistic pathogens, but have also been isolated from a wide variety of environmental sources (Reverdy et al., 1984; Rihs et al., 1993). Various *Acinetobacter* and *Streptococcus* species have physiological functions in their mammalian hosts, but the genera also include pathogenic and/or environmental species (Krzyściak et al., 2013; Touchon et al., 2014). *Fusobacteria* species are typically closely related to mucous membranes, especially the oral cavities, while *Cutibacterium* and *Kocuria* species have most often been isolated from the skin (Hofstad, 1992; Savini et al., 2010; Scholz and Kilian, 2016). One of our negative controls yielded *Kocuria* colonies. This was the only taxon shared between the actual meconium samples and the negative controls, suggesting low-level contamination from the sampling environment.

Although we detected cultivable bacteria in some of the meconium samples, most of the samples were culture negative. This indicates that the presence of live bacteria in the fetal intestine is rare, in contrast to the prevalent bacterial DNA. Individual differences in amounts of viable bacteria retrieved from meconium were also described in a recent dog study. Interestingly, puppies born without detectable meconium microbiota were shown to have a slower growth rate than those in which meconium microbiota were detected (Pipan et al., 2020). Both the mucosal surface of the intestine as well as the amniotic fluid contain bacteriostatic substances, such as lactoferrin and salivary scavenger and agglutinin (SALSA), which are able to suppress bacterial growth and viability (Reichhardt and Meri, 2016; Lisowska-Myjak et al., 2019). In human meconium, SALSA amounts to 10 % of all proteins, highlighting its potential role in antimicrobial defense (Reichhardt et al., 2014). Lactoferrin, SALSA and other similar substances may explain the small number of viable bacteria observed in our study, in contrast to the significant amount of detected bacterial DNA.

The culture-independent techniques were performed based on the 16S rRNA gene. Since contamination is a major challenge in the analysis of low bacterial biomass samples, we processed several types of negative controls alongside our biological samples, treated qPCR mastermix with dsDNAse, and applied rigorous *in silico* filtering of potential contaminants (Glassing et al., 2016; Eisenhofer et al., 2018). Many similar studies are published without or with a very limited number of contamination controls, which makes our approach more reliable, especially for assessing low bacterial biomass environments.

We observed a small but significant amount of bacterial DNA in the meconium samples by 16S qPCR, suggesting a prenatal transmission of bacterial DNA to the fetal intestine. This is consistent with previous studies in cattle, in which calves born at term *per vias naturales*, were sampled (Mayer et al., 2012; Alipour et al., 2018; Klein-Jöbstl et al., 2019). A profile dominated by the phyla Proteobacteria, Firmicutes, Bacteroidetes and Actinobacteria was identified in the meconium, in agreement with previous studies using 16S rRNA gene amplicon sequencing (Alipour et al., 2018; Yeoman et al., 2018; Klein-Jöbstl et al., 2019; Husso et al., 2020; Guzman et al., 2020). Additionally, some of the most prevalent genera observed in meconium in our study, such as *Sphingomonas*, *Acinetobacter*, *Pseudomonas*, *Ruminococcus* and *Stenotrophomonas,* have also been previously described in neonatal calves (Klein-Jöbstl et al., 2019).

In contrast to the meconium, there was no significant difference between the 16S rRNA gene copy numbers of amniotic fluid samples and the negative controls, and all were culture negative. A recent caesarean section study in sheep stated amniotic fluid sterility during the third trimester of pregnancy (Malmuthuge and Griebel, 2018). In other studies however, a low biomass microbiome in the pregnant bovine uterus was described (Moore et al., 2017; Guzman et al., 2020).

Using 16S rRNA gene amplicon sequencing, we found a bacterial signature in the amniotic fluid, dominated by the phyla Proteobacteria, Firmicutes, Bacteroidetes and Actinobacteria, in accordance with previous studies (Karstrup et al., 2017; Moore et al., 2017; Guzman et al., 2020). Deeper taxonomic identification, down to the family or genus level, revealed no further agreements with these studies.

From a comparative point of view between mammalian species, the amniotic fluid sampled by Collado et al. (2016) in women during pre-labor C-section was found to contain a distinct, low diversity, low biomass microbiome, predominated by Proteobacteria (*Enterobacteriaceae*). Despite this and other recent studies, the presence of microbial DNA and culturable bacteria in the human placenta and fetal environment is still under debate (de Goffau et al., 2019).

The microbial DNA signatures in meconium and amniotic fluid were similar at the phylum level. However, they presented a significantly different microbial composition at genus and ASV level: genera *Staphylococcus*, *Rubrobacter* and *Clostridium* were relatively more abundant in meconium samples than in amniotic fluid samples according to DESeq2 analysis. Complementing this with the LefSe analysis using all taxonomic levels, also found *Clostridiales* and *Rubrobacter* taxons relatively more abundant in the meconium. The microbial compositions in the bovine amniotic fluid and meconium of the corresponding neonate did not show a significant correlation, contrary to earlier studies in humans (Collado et al., 2016; He et al., 2020).

Our analysis suggests that the bacterial DNA found in meconium does not necessarily originate from the amniotic fluid. The maternal microbial components may be translocated via the placental and umbilical blood vessels directly to the fetal internal organs. Microbial macromolecules and even live bacteria may be transported by leukocytes via these blood vessels (Perez et al., 2007).

A recent bovine study revealed differing microbial communities between the amniotic fluid and the meconium (Guzman et al., 2020). As the fetus swallows the amniotic fluid, which is concentrated and retained in the intestine as meconium, together with epithelial cells and intestinal secretions, it is unexpected that the two would harbor completely different bacteria. However, the amniotic fluid microbiota may fluctuate dynamically over time, while meconium represents a more stable collection of substances accumulated over the gestation period. Moreover, fetal excretion in the amniotic fluid is limited in cattle. Meconium is usually expelled only after birth, and fetal urine largely accumulates in the allantoic cavity, between chorion and amnion, rather than in the amniotic cavity (Bongso and Basrur, 1976).

The less permeable synepitheliochorial placenta evidenced by three distinct layers may restrict the translocation of bacteria and their components from the dam to the fetus in cattle, in comparison to humans. The very low levels of bacterial DNA in the bovine amniotic fluid limit the sequencing depth and thus the precision of the compositional analyses. Our strict data decontamination protocol furthermore minimized the artefactual similarity arising from shared contaminants.

In conclusion, we found that the meconium of full-term calves delivered by elective caesarean section in most cases contains a small amount of diverse bacterial DNA and eventually although rather rare, culturable bacteria. In the amniotic fluid, bacteria were not observed by 16S qPCR or culturing, but a microbial DNA profile was distinguishable from controls by amplicon sequencing. Based on these results, bacterial components are translocated to the fetus *in utero*, but the prenatal acquisition of live bacteria is likely not physiologically significant.

## Data availability

The raw 16S rRNA gene amplicon sequencing dataset is fully available in the NCBI Sequence Read Archive (SRA) with the accession number PRJNA643145 after journal acceptance.

## Acknowledgements

We thank the owners of the cows for allowing sampling of their animals, and all clinicians and graduate veterinary students for helping with the monitoring of the animals, performing the surgery and sampling. Additionally, we thank Kirsi Lahti for expert technical assistance in the laboratory.

## Author contributions

MN, LL, AI and GO contributed to conception and design of the study. AH performed the bioinformatics, statistical analyses and bacterial culture. AH and TG performed MALDI-TOF analyses. LL and JG organized the sampling and collected the samples. AH and LL wrote the manuscript. All authors contributed to manuscript preparation, read, and approved the submitted version.

## Ethics declaration

All experimental procedures were approved by the institutional ethics and animal welfare committee of the Faculty of Veterinary Medicine (EC2018/002 - Ghent University, Belgium). The cows’ owners were informed about the study and gave their written consent for their cows’ participation.

