## Supplemental info for "The composition of the microbiota in the full-term fetal gut and amniotic fluid: a bovine caesarean section study"

**Supervised the study

#### Supplementary results

**Supplementary Table 1. Most prevalent and abundant genera in the meconium and amniotic fluid samples.**

| *Taxon* | *Mec-Preva* | *Mec-Ave* | *Mec-SD* | *Amn-Preva* | *Amn-Ave* | *Amn-SD* |
|---|---|---|---|---|---|---|
| **Actinobacteria** | **100 %** | **9.3 %** | **5.3 %** | **100 %** | **9.3 %** | **5.7 %** |
| Corynebacterium 1 | 76 % | 1.5 % | 2.0 % | 67 % | 1.1 % | 1.2 % |
| Rubrobacter | 71 % | 1.3 % | 1.4 % | 17 % | 0.1 % | 0.2 % |
| Cutibacterium | 35 % | 0.4 % | 0.8 % | 50 % | 1.2 % | 1.7 % |
| Kocuria | 47 % | 0.5 % | 0.8 % | 29 % | 0.5 % | 1.0 % |
| Mycobacterium | 47 % | 0.3 % | 0.5 % | 17 % | 0.2 % | 0.7 % |
| Nocardioides | 47 % | 0.5 % | 0.8 % | 33 % | 0.3 % | 0.5 % |
| Lawsonella | 0 % | 0.0 % | 0.0 % | 46 % | 1.6 % | 3.1 % |
| Micrococcus | 41 % | 0.6 % | 1.2 % | 29 % | 0.5 % | 1.3 % |
| **Bacteroidetes** | **100 %** | **12.3 %** | **7.9 %** | **96 %** | **10.2 %** | **8.5 %** |
| Bacteroides | 53 % | 2.1 % | 3.1 % | 25 % | 0.9 % | 2.8 % |
| Chryseobacterium | 53 % | 0.6 % | 0.9 % | 46 % | 1.4 % | 2.4 % |
| Cloacibacterium | 29 % | 1.3 % | 4.4 % | 42 % | 2.0 % | 5.3 % |
| Alistipes | 41 % | 0.5 % | 0.6 % | 0 % | 0.0 % | 0.0 % |
| Prevotellaceae UCG-003 | 41 % | 0.6 % | 1.0 % | 0 % | 0.0 % | 0.0 % |
| Alloprevotella | 35 % | 0.5 % | 0.7 % | 13 % | 0.3 % | 0.8 % |
| Rikenellaceae RC9 gut group | 35 % | 0.8 % | 1.4 % | 21 % | 0.2 % | 0.6 % |

| | | | | | | |
|---|---|---|---|---|---|---|
| **Firmicutes** | **100 %** | **30.0 %** | **16.5 %** | **100 %** | **34.0 %** | **16.2 %** |
| Streptococcus | 59 % | 2.5 % | 6.1 % | 88 % | 6.0 % | 5.5 % |
| Staphylococcus | 76 % | 4.2 % | 4.6 % | 83 % | 6.4 % | 5.8 % |
| Clostridium sensu stricto 1 | 71 % | 3.7 % | 8.6 % | 29 % | 0.7 % | 1.4 % |
| Enterococcus | 41 % | 0.5 % | 0.7 % | 63 % | 4.8 % | 9.5 % |
| Bacillus | 59 % | 1.1 % | 1.5 % | 54 % | 3.9 % | 12.9 % |
| unclassified Lachnospiraceae | 59 % | 1.2 % | 1.4 % | 13 % | 0.2 % | 0.7 % |
| Christensenellaceae R-7 group | 53 % | 1.3 % | 1.9 % | 13 % | 0.1 % | 0.2 % |
| Lactobacillus | 29 % | 0.3 % | 0.9 % | 46 % | 1.8 % | 3.2 % |
| Ruminococcaceae UCG-010 | 41 % | 0.5 % | 0.7 % | 4 % | 0.2 % | 0.8 % |
| Ruminococcaceae UCG-013 | 41 % | 0.7 % | 1.1 % | 17 % | 0.5 % | 1.7 % |
| Ruminococcaceae UCG-014 | 41 % | 0.6 % | 0.9 % | 8 % | 0.1 % | 0.2 % |
| [Ruminococcus] gauvreauii group | 35 % | 0.3 % | 0.5 % | 8 % | 0.1 % | 0.4 % |
| Acetitomaculum | 35 % | 0.3 % | 0.5 % | 4 % | 0.0 % | 0.1 % |
| Aerococcus | 35 % | 0.4 % | 0.8 % | 13 % | 0.1 % | 0.4 % |
| Family XIII AD3011 group | 35 % | 0.3 % | 0.5 % | 13 % | 0.0 % | 0.1 % |
| Mogibacterium | 35 % | 0.5 % | 0.7 % | 17 % | 0.4 % | 1.1 % |
| Ruminococcaceae UCG-005 | 35 % | 0.6 % | 1.2 % | 21 % | 0.5 % | 1.2 % |
| Turicibacter | 35 % | 0.4 % | 0.6 % | 13 % | 0.2 % | 0.8 % |
| unclassified Peptostreptococcaceae | 35 % | 0.7 % | 1.2 % | 13 % | 0.1 % | 0.2 % |
| unclassified Ruminococcaceae | 35 % | 0.3 % | 0.6 % | 4 % | 0.0 % | 0.2 % |
| **Proteobacteria** | **100 %** | **44.4 %** | **18.9 %** | **100 %** | **43.0 %** | **14.8 %** |
| Delftia | 94 % | 11.8 % | 10.8 % | 92 % | 5.0 % | 3.6 % |
| Acinetobacter | 88 % | 2.7 % | 2.3 % | 71 % | 3.1 % | 3.0 % |
| unclassified Burkholderiaceae | 82 % | 2.1 % | 2.5 % | 58 % | 1.1 % | 1.6 % |
| Sphingomonas | 47 % | 0.9 % | 1.5 % | 71 % | 5.0 % | 6.3 % |
| Brevundimonas | 71 % | 0.9 % | 1.2 % | 38 % | 0.6 % | 1.4 % |
| Paracoccus | 47 % | 1.6 % | 3.7 % | 63 % | 2.0 % | 3.2 % |
| Massilia | 59 % | 2.8 % | 7.4 % | 29 % | 0.5 % | 1.1 % |
| Methylobacterium | 59 % | 0.9 % | 1.0 % | 46 % | 0.7 % | 1.1 % |
| Enhydrobacter | 53 % | 0.7 % | 1.0 % | 0 % | 0.0 % | 0.0 % |
| Pseudomonas | 47 % | 2.5 % | 8.2 % | 50 % | 1.7 % | 2.2 % |
| Psychrobacter | 47 % | 0.7 % | 1.1 % | 25 % | 0.9 % | 2.2 % |
| Stenotrophomonas | 47 % | 0.6 % | 0.9 % | 42 % | 0.8 % | 1.4 % |
| Cupriavidus | 41 % | 0.4 % | 0.6 % | 8 % | 0.1 % | 0.3 % |
| Novosphingobium | 41 % | 0.7 % | 1.2 % | 17 % | 0.1 % | 0.3 % |
| Alkanindiges | 35 % | 0.6 % | 1.2 % | 8 % | 0.1 % | 0.2 % |
| Allorhizobium-Neorhizobium-Pararhizobium-Rhizobium | 35 % | 0.5 % | 0.9 % | 17 % | 0.2 % | 0.7 % |
| Haemophilus | 35 % | 0.5 % | 0.7 % | 17 % | 0.6 % | 1.7 % |
| Rubellimicrobium | 35 % | 0.6 % | 2.0 % | 33 % | 0.6 % | 1.2 % |
| Sphingopyxis | 35 % | 0.3 % | 0.8 % | 21 % | 0.2 % | 0.5 % |
| Neisseria | 29 % | 0.4 % | 1.0 % | 33 % | 0.7 % | 1.4 % |

### Supplementary methods

#### *Detailed description of the bioinformatics pipeline*

##### Obtaining the data

The sequencing data for 69 samples was obtained from the sequencing lab in demultiplexed FASTQ format. The compressed .tar.gz-package was uploaded to the now decommissioned supercluster *Taito* of Finnish Center for Scientific Computing (CSC). Package md5sum was checked before further decompression and analyses.

##### Quality check

FastQC 0.11.8 was ran for all the files and the resulting reports were checked manually[1]. FastQC reports were also compiled and assessed with MultiQC 1.7[2].

##### Trimming

All leftover primers and spacers were removed with Cutadapt v.1.10[3].

find -name "*R1_001.fastq.gz" -exec cutadapt -g CCTACGGGNGGCWGCAG -o '{}.TRIMMED_CUTADAPT_FW.gz' '{}' ';'

find -name "*R2_001.fastq.gz" -exec cutadapt -g GACTACHVGGGTATCTAATCC -o '{}.TRIMMED_CUTADAPT_REV.gz' '{}' ';'

The trimmed sequences were again checked with FastQC 0.11.8 and MultiQC 1.7[1,2].

##### Mapping file

A QIIME 2 compatible mapping file containing the sample metadata was created with Google Sheets and validated with Keemei[4,5].

##### QIIME2

Fastq.gz files were first imported to QIIME2 v2019.4 with command qiime tools import. The sequences were checked again using command qiime demux summarize[5]. The mean number of raw sequences was 114380 (max 187955 , min 31, total 7892241) with read quality starting to drop below 20 in forward direction at 280 and reverse at 220. DADA2 was ran using the command dada2 denoise-paired with truncating option F280 and R220[10]. This results in an amplicon sequence variant (ASV) table.

Stats from the denoising and merging steps were checked with qiime metadata tabulate. Resulting data was explored with visual summaries created with qiime feature-table summarize and qiime feature-table tabulate-seqs.

The phylogenetic tree was created using qiime alignment mafft[6]. Highly variable positions adding noise were masked with qiime alignment mask. FastTree was used to create a phylogenetic tree from the masked alignment: qiime phylogeny fasttree and qiime phylogeny midpoint-root[7].

Preliminary diversity analysis was created with qiime diversity core-metrics-phylogenetic using sampling depth 15988.

The taxonomy was assigned according to the SILVA v132 QIIME release 99 %[8]. Representative set of sequences and taxonomy were imported to QIIME2 with qiime tools import. Reference reads were first extracted from the sequence directory with qiime feature-classifier extract-reads[9]. Then a Naïve Bayes classifier was trained to the curated taxonomy with qiime feature-classifier fit-classifier-naïve-bayes[10]. Finally the actual classification was called with qiime feature-classifier classify-sklearn[11].

The taxonomy was visualised again with qiime metadata tabulate and qiime taxa barplot. After this the ASV table was exported from QIIME2 for handling in spreadsheet programs. First the table and taxonomy data was exported in biom-format with qiime tools export. The two were combined with biom package: biom add-metadata[13]. Finally the ASV table and table with taxonomy were converted to .tsv format with biom convert.

#### Data decontamination

Mitochondria, chloroplasts and singletons were filtered out, leaving only bacteria with at least phylum-level identification.

The processed data was filtered further to remove ASVs which represented probable contaminants[12]. An ASV was removed if its prevalence in actual samples was ≤2× its prevalence in instrument controls

(DNA extracted from empty sampling instruments only handled aseptically in a laminar flood cabinet) or its prevalence in field controls (empty sampling instruments exposed to the surgery room environment), and if its mean relative abundance in actual samples was ≤10× its mean abundance in instrument controls or its mean abundance in field controls. The filtering was performed separately for meconium and amnion samples. If less than 500 reads remained after the decontamination, as in six meconium samples, the sample was removed from further analyses.
